# Embigin deficiency leads to delayed embryonic lung development and high neonatal mortality

**DOI:** 10.1101/2021.07.05.451131

**Authors:** Salli Talvi, Johanna Jokinen, Kalle Sipilä, Pekka Rappu, Fu-Ping Zhang, Matti Poutanen, Pia Rantakari, Jyrki Heino

## Abstract

Embigin (gp70), a transmembrane glycoprotein, has been shown to regulate hematopoietic stem cell and progenitor cell niche. Still, little is known about its expression and function in other organ systems during development or adulthood. By combining immunofluorescence, RNA sequencing, and *in vivo* mouse models, we show that embigin is highly expressed during development and in adult lung, kidney, epididymis, skin, and testis. Adult Emb^-/-^ mice have a normal lifespan and fertility without apparent pathologies. In contrast, the Emb^-/-^ embryos are significantly smaller than their WT littermates. Markedly increased mortality of the Emb^-/-^ embryos is seen especially during the neonatal period. Embigin is present in the placenta, but placental morphology and gene expression patterns stay unaltered. At E17.5, Emb^-/-^ mice show defective morphogenesis of the lung, low alkaline phosphatase activity in amniotic fluid, and remarkable activation of genes involved in cell proliferation in the lungs. Thus, lung underdevelopment explains the high neonatal mortality. Our work demonstrates the crucial role of embigin during development, and it paves the way to further characterization of embigin in specific organ systems in development and homeostasis.

**Summary statement:** Embigin is a basigin-group transmembrane glycoprotein. *In vivo* mouse model shows that embigin is crucial for embryonic lung development and neonatal survival.

## Introduction

Embigin (gp70) is a highly glycosylated member of the basigin subgroup that belongs to the immunoglobulin superfamily (Huang et al., 1993;Ozawa et al., 1988). In addition to embigin, the group includes two other type I membrane proteins, namely basigin (EMMPRIN/CD147) and neuroplastin (Np65/gp65 and Np55/gp55). These three proteins have evolutionarily conserved domains, and they are proposed to be involved in similar cellular functions, including the regulation of cell adhesion, migration, and metabolism (Muramatsu and Miyauchi, 2003;Williams and Barclay, 1988). In this study, we will focus on embigin, the least known member of the basigin group.

Structurally, all three members of basigin subgroup resemble each other, having an amino acid sequence identity of 37% - 46% (Hanna et al., 2003). They share the overall structure possessing an extracellular immunoglobulin-like (Ig-like) domain, a single hydrophobic transmembrane domain, and a short cytoplasmic tail. However, there are also significant structural differences that most probably contribute to the biological roles of the proteins, for example, both basigin-1 and neuroplastin Np65 are composed of three Ig-like domains, while basigin-2, embigin, and neuroplastin Np55 have only two. The other two basigin isoforms found in human, basigin-3 and basigin-4, are structurally small and they comprise only one Ig-like domain (Liao, C. et al., 2011). In addition to the variability in the number and the sequence of the Ig-like domains, the N-glycosylation state of these highly glycosylated proteins may define the function of the protein (Langnaese et al., 1997;Ochrietor et al., 2003;Yoshida et al., 2000;Yu et al., 2008).

Now, more than 30 years after the first discovery of embigin, knowledge of the protein expression patterns in both mouse and human tissues is still dispersed and partly incoherent. As an example, the embigin protein expression pattern is not available through the Human Protein Atlas because the existing data provide inconclusive results. However, strong embigin mRNA expression has been localized to mouse embryos during the early phases of the development (Fan et al., 1998;Huang et al., 1990). In adult mice and rats, only low levels of embigin mRNA have been reported in several organs (Guenette et al., 1997;Huang et al., 1990). While other basigin group members have been detected to display a multifunctional nature, the biological role of embigin is not understood yet. For example, basigin has been reported to act in a wide variety of cellular processes including development, activation, proliferation, migration, invasion, and adhesion in T lymphocytes (Hahn et al., 2015). Also neuroplastin, which is enriched in neurons and synapses, has basigin-like functions but in more restricted locations (Beesley et al., 2014;Hill et al., 1988;Langnaese et al., 1997;Smalla et al., 2000). Given the variety of cellular functions that the basigin family members are involved in, it is not surprising that they also have connections to pathological processes, such as cancer (Nabeshima et al., 2006;Riethdorf et al., 2006). To date, embigin has been reported to be a suppressor of tumorigenesis in breast cancer (Chao et al., 2015) and a promoter of epithelial-mesenchymal transition in pancreatic carcinoma (Jung et al., 2016).

All three members of the basigin group are involved in cellular metabolism (Kirk et al., 2000). They have been reported to escort monocarboxylate transporters (MCTs), the carriers of molecules such as L-lactate and pyruvate, to the plasma membrane (Fisel et al., 2018). Embigin is identified as a primary ancillary protein for MCT2 (Wilson et al., 2005), but it might act as the assisting protein also for MCTs 1, 3 and 4 (Halestrap, 2013;Skiba et al., 2021). A recent paper has shed more light on the complex mechanism of the MCT function and unveiled a direct interaction between basigin or embigin and carbonic anhydrase IV (CA IV) (Forero-Quintero et al., 2019). CA IV is a metalloenzyme that also facilitates the transport activities of specific MCTs (Becker et al., 2005;Becker et al., 2010;Klier et al., 2011). Besides the potential role of embigin in the MCT and CA IV translocation, only a few embigin interaction partners have been reported. It has been suggested that embigin may regulate cell adhesion by modifying the integrin function (Huang et al., 1993). Embigin has also been reported to interact with galectin-3 (Dange et al., 2017) and S100A4 protein (Ruma et al., 2018). Furthermore, embigin has been identified as a bone marrow stem cell niche factor, more specifically as a hematopoietic stem/progenitor cell quiescence regulator (Silberstein et al., 2016). During the maturation of bone marrow progenitor cells, embigin seems to be specifically repressed by a transcription factor Pax5 in B lymphocytes (Pridans et al., 2008). In addition to the putative participation in these processes, the physiological role of embigin is still poorly understood.

Here, we unveil the expression pattern of embigin protein during mice embryonic development and in adult mice. Besides, we shed light on the biological function of embigin during development. Our embigin knockout mouse model and RNA sequencing of embryonic lungs confirm that embigin is a critical protein for overall embryonic growth, particularly for early lung development.

## Results

### Embigin is expressed from early mouse embryonic development into adulthood

The knowledge of embigin expression in mouse tissues is dispersed and partly incoherent. Therefore, we used the whole-mount immunofluorescence technique to visualize the embigin protein expression pattern in mice embryos at embryonic days E8.5 - E10.5. In agreement with previous studies (Fan et al., 1998;Huang et al., 1990), the most robust embigin expression was detected at the early stage of embryogenesis, and embigin was observed to be an abundant protein specifically in the developing gut (Fig. 1A). However, embigin expression did not cease after E10.5, albeit an apparent decrease in its expression was observed. At E13.5, low embigin expression was detected in restricted tissues such as kidney, lung, and small intestine (Fig. S1A), and later in gestation, at E17.5, increased embigin expression was observed in the kidneys (Fig. 1B). Thus, unlike stated in the previous reports, embigin expression is not restricted to the early embryonic development of the mouse but continues throughout gestation.

**Figure 1.**
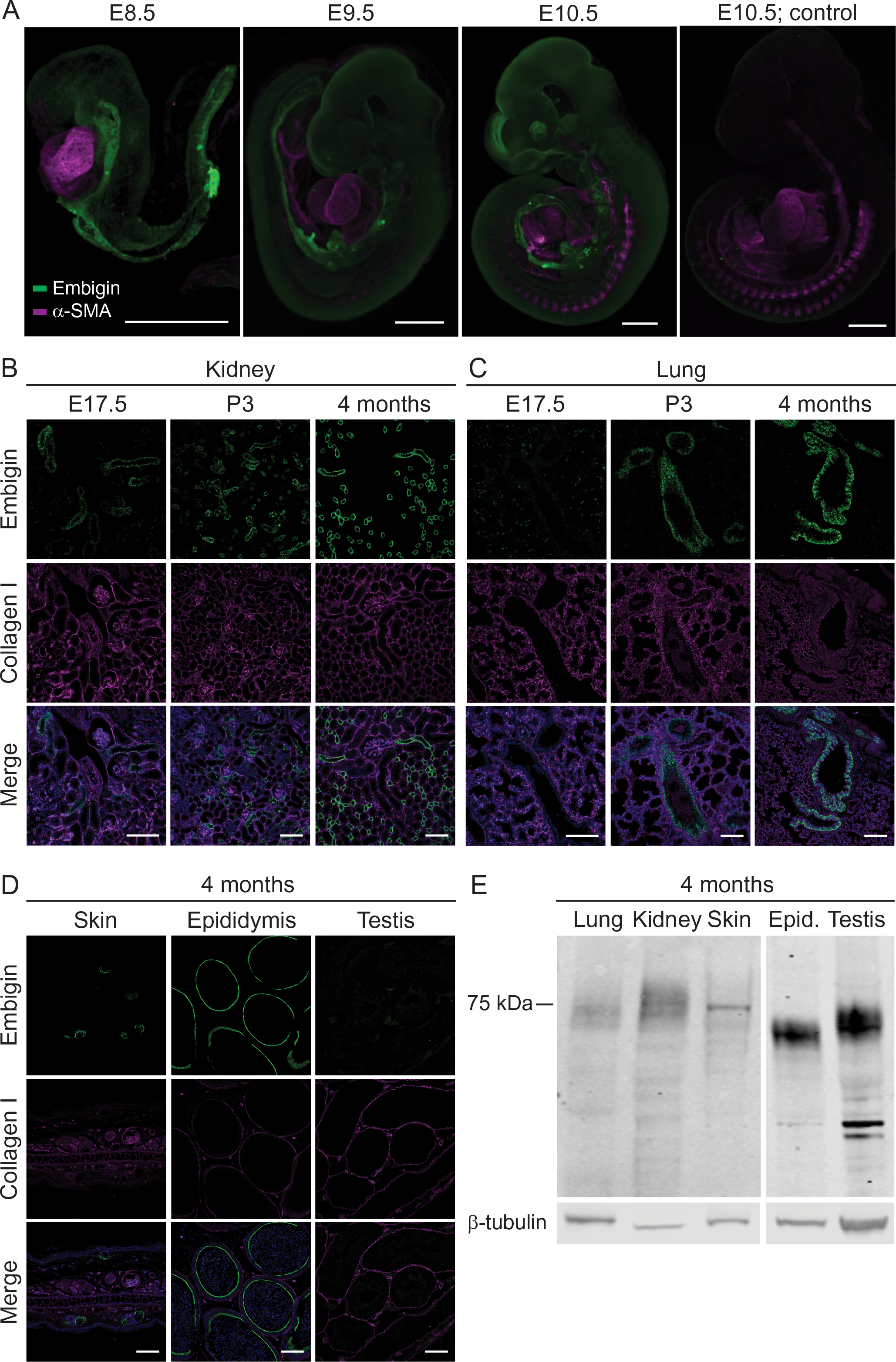
Embigin is widely expressed until E10.5 and after that in specific organs. (A) WT mouse embryos at embryonic days E8.5, E9.5 and E10.5 were stained with embigin and α-smooth muscle actin (α-SMA) antibodies by using the whole-mount immunostaining technique. As the negative control, secondary antibody only was used; α-SMA was stained as a positive control. Representative images are shown. Scale bars: 500 µm. (B – D) The paraffin sections of the kidney (B), lung (C), skin, epididymis, and testis (D) were immunostained with embigin and collagen I antibodies. Collagen I was stained as a positive control. Samples were collected at E17.5 (B, C), postnatal day P3 (B, C), and four-month-old mice (B, C, D). Scale bars: 100 µm. (E) Embigin expression in protein samples extracted from four-month-old WT mouse lung, kidney, skin, epididymis, and testis tissues were studied by Western blotting. β-tubulin was used as a control. The blot image was cut to show only embigin positive organs.

We also determined the precise location of embigin after the gestational period. A set of adult organs at the age of four months was analyzed using immunofluorescence microscopy. Kidney (Fig. 1B), lung (Fig. 1C), epididymis, and skin (Fig. 1D) showed high embigin levels. In the kidneys, embigin was located in the epithelial cells lining tubular structures (Fig. 1B), and also the epithelial cells of the lung airways were shown to be highly embigin positive (Fig. 1C). Furthermore, the tubular structures in the caudal pole of the epididymis were determined to have a strong embigin expression, whereas, in the skin, embigin expression was located in the sebaceous glands (Fig. 1D). In both lungs and kidneys, the expression level of embigin was shown to elevate shortly after birth at P3, and the expression was observed to be the strongest in the adult mice (Figs 1B, C). No differences between male and female mice were observed, apart from the embigin expression in the male epididymis.

The embigin protein expression in the organs described above was also further confirmed by Western blot (Fig. 1E). A strong signal was also detected in the testis suggesting the presence of embigin in the tissue. Heart, liver, spleen, small intestine, adrenal glands, and ovary were determined as embigin negative tissues (Fig. S1B). In the embigin positive tissues, embigin was detected as a characteristic broad protein band ranging from about 60 to 90 kDa. The observed variation of the approximated molecular mass is typical for embigin and can be explained by the differential glycosylation of the nine potential glycosylation sites in the protein (Ozawa et al., 1988). The highest molecular mass of embigin was observed in the kidney, while the lowest molecular mass was found in the epididymis, and the smallest variation was detected in the skin (Fig. 1E). Thus, the degree of embigin glycosylation was shown to vary in a tissue-dependent manner, which might implicate the distinct function of embigin in these tissues. Together, our results confirm that the embigin protein is expressed in the specific structures of the lung, kidney, skin, epididymis, and testis of the four-month-old mice.

### Embigin deficiency leads to an increase in neonatal mortality

To examine the role of embigin *in vivo*, knockout mice lacking the exon 5 of the embigin gene were generated (Fig. S2A). The absence of embigin expression in the embigin deficient (Emb^-/-^) mice was confirmed both by PCR (Fig. S2B) and by performing Western blot analysis of the kidney tissues (Fig. S2C). In the Western blot analysis, a typical diffuse band around 75 kDa was observed only in the kidneys of the wild type (WT) mice (Fig. S2C). Furthermore, final verification of the lack of embigin in Emb^-/-^ mice was gained through staining Emb^-/-^ and WT E9.5 embryos with embigin and alpha smooth-muscle actin (α-SMA) antibodies (Fig. S2D).

Using Emb^-/-^ mice, the effect of the embigin deficiency on the lifespan of affected animals was studied next. The genotypes of 203 embryos from Emb^+/-^ heterozygous intercrosses at the ages between E8.5 and E17.5 were analyzed. At embryonic day E8.5, 25% of all embryos were embigin deficient. Thus, the relative frequency of the genotypes at this embryonic stage was found to follow the Mendelian distribution. When embryos at E17.5 were inspected, the frequency of Emb^-/-^ embryos was found to be only 18% instead of the expected 25% (Fig. 2A). Spearman’s rank correlation analysis indicated that the small gradual decrease in the number of embigin null embryos as the gestation progressed was statistically significant (r_s_ = -0.837, p = 0.019; Fig. 2A). The frequency of Emb^+/-^ embryos was also decreasing, but not in a statistically significant magnitude (r_s_ = -0.667, p = 0.102; Fig. 2A). Next, the genotypes of 284 pups from 40 different litters were examined between P14 and P21. Based on the study, only 7% of the pups from Emb^+/-^ breedings were embigin deficient at P14-P21 (Fig. 2B). 42% survived Emb^-/-^ pups were males and 58% females. Our results indicate that based on the Mendelian expectation, 28% of Emb^-/-^ pups were lost already in the prenatal period, and in total, 72% of expected Emb^-/-^ pups did not reach adulthood.

**Figure 2.**
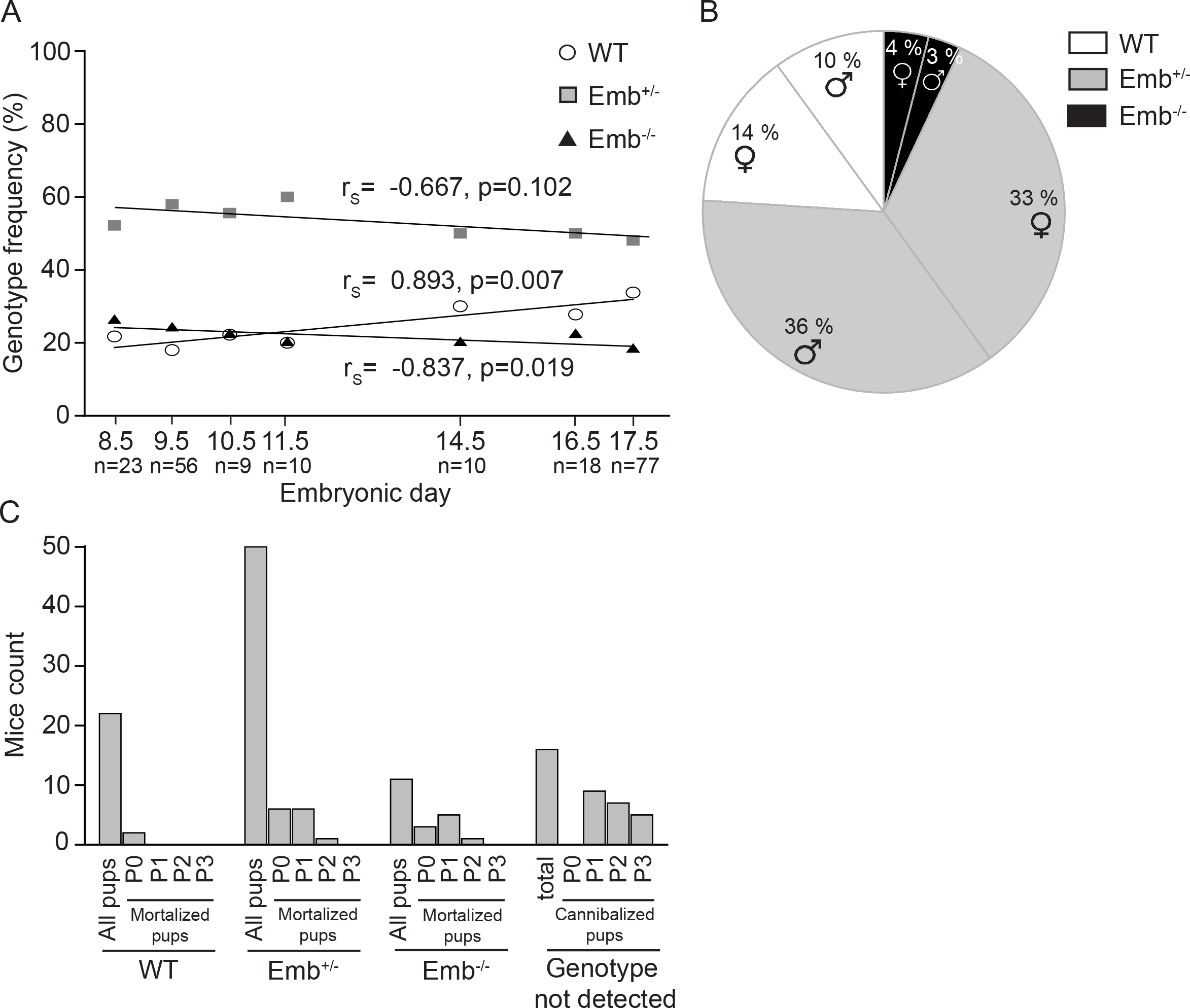
Embigin deficiency increases mortality from embryonic day E8.5 to P3. (A) The relative frequency of the genotypes of the pups from heterozygous Emb^+/-^ breedings were determined. Three litters at E8.5; 6 litters at E9.5; 1 litter at E10.5, E11.5, and E14.5; 2 litters at E16.5, and 10 litters at E17.5 were examined (n = number of pups analyzed). Correlation between the frequency of each genotype and the embryonic stage was analyzed by calculating Spearman’s rank correlation coefficient (r_s_) and its statistical significance. (B) The relative frequency of the genotypes of the pups from Emb^+/-^ breedings were determined. 284 pups from 40 litters were analyzed at P14-P21. (C) To determinate the postnatal survival frequency of the Emb^-/-^ mice, 100 pups from 17 litters and 6 different Emb^+/-^ breedings were followed after birth. The genotypes were determined after the death of the pup.

To determine the time point at which the lethality of Emb^-/-^ mice occurred, 100 pups from 17 litters from six different Emb^+/-^ breedings were followed up after birth. These pups were genotyped after their death. The results indicate that most of the Emb^-/-^ pups died during days 0 and 1 in postnatal life (Fig. 2C). Noteworthy, the mortality of Emb^+/-^ mice also appeared to be slightly elevated. While 81% of the born Emb^-/-^ mice died during the first three postnatal days, 26% of Emb^+/-^ and only 9% of WT mice were lost during the period. Furthermore, 16% of the pups were fully cannibalized during postnatal days of P1-P3 before they were genotyped. While the frequency of Emb^-/-^ embryos slightly decreased already before birth, the results indicate that the major loss of Emb^-/-^ mice occurred during the neonatal period. It cannot be excluded, however, that some pups were lost already during the parturition.

### Embigin deficiency does not affect the lifespan of the mice after the neonatal period

To study the effect of embigin deficiency on mice that survived beyond the first three postnatal days, Emb^-/-^ and WT mice were further analyzed at the age of 2, 4, or 6 months. The obtained results indicated that there was no difference in body weights when the adult WT and Emb^-/-^ mice were compared (Fig. S3A). Furthermore, neither the histology nor weights of specific organs were different (Figs S3B, C, D; Table S1). To assess whether embigin deficiency could affect the fertility of mice, ten pairs of Emb^-/-^ mice were allowed to breed. Four out of ten breedings did not produce viable pups and out of the 23 litters produced, ten were fully cannibalized. On average 2.3 pups per litter survived and reached adulthood (Table 1). These observations were consistent with the high mortality rate of embigin deficient embryos and newly born mice. Based on the data, the embigin deficient mice that survive are fertile, and they have changes neither in the typical body and organ weights nor the histological architecture of the tissues studied.

**Table 1.** Emb^-/-^ mice are fertile. Ten Emb^-/-^ homozygous breedings were followed until the pups were genotyped at P14-P21.

|  |  |
| --- | --- |
| Emb<sup>-/-</sup> breeding pairs | 10 |
| Emb<sup>-/-</sup> breeding pairs with survived litters | 6 |
| Cannibalized litters | 10 |
| Litters with survived pups | 13 |
| Survived female pups | 14 |
| Survived male pups | 16 |
| Average litter size | 2.3 |

### Embigin deficiency causes delayed growth of mouse embryos

While the weights of four-month-old Emb^-/-^ mice did not differ from WT mice, the body sizes of Emb^-/-^ embryos tend to be smaller than their Emb^+/-^ or WT littermates as imaged at E11.5, E14.5, and E17.5 in Fig. 3A. Furthermore, the body weights of the Emb^-/-^ embryos at E17.5 were significantly (p = 0.001) smaller than their littermates, average body weights being 690 ± 54 mg for Emb^-/-^ mice and, 916 ± 136 mg for Emb^+/-^ and 834 ± 134 mg for WT embryos (Fig. 3B). The fact that the normal function of the placenta is pivotal for optimal fetal growth and development led us to characterize the placentas of Emb^-/-^ and WT embryos from Emb^+/-^ breedings. Between E11.5 and E17.5, an increasing embigin expression was detected in the labyrinthine layer of the placenta in WT embryos, but not in Emb^-/-^ embryos, by using immunofluorescence staining technique (Fig. 3C). Though the intensive embigin expression was detected in the placenta, histological differences were not observed between the placentas of WT and Emb^-/-^ embryos (Fig. 3D). Furthermore, based on the RNA sequencing data, only four genes, one of them being embigin, were differentially expressed in the placentas of five Emb^-/-^ embryos compared to the placentas of five WT embryos at E17.5 (Fig. 3E). The gene was determined to be differentially expressed only if log2 of fold change value was above 0.6 or below -0.6 and Benjamini-Hochberg-corrected p-value less than 0.05. Taken together, these results do not support the idea that embigin deficiency could cause placental dysfunction that would manifest as a fetal growth restriction observed in Emb^-/-^ mice. Therefore, other vital organs were examined next.

**Figure 3.**
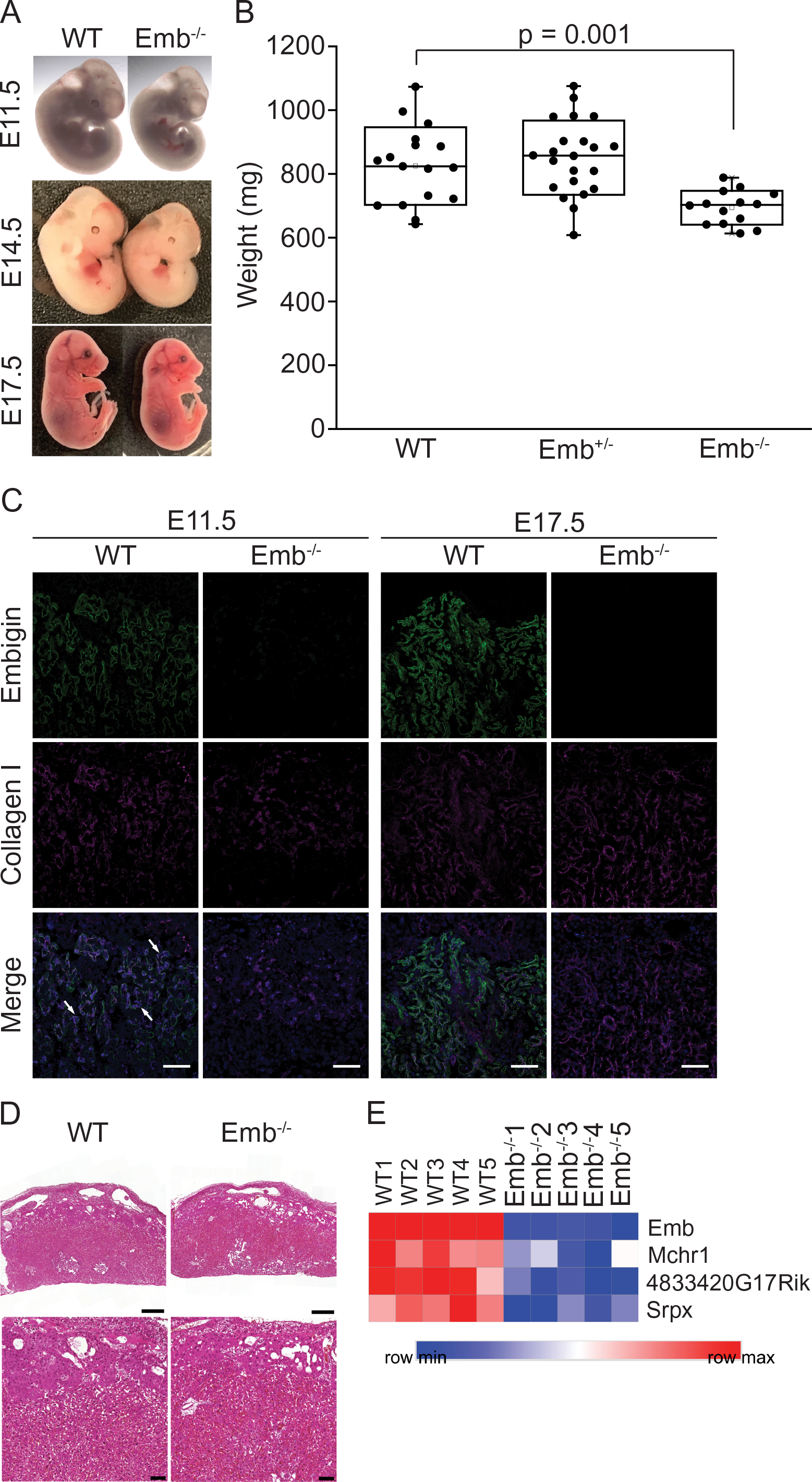
The smaller size of Emb^-/-^ embryos is not caused by a placental failure. (A) Representative images of WT and Emb^-/-^ mice are shown at E11.5, E14.5, and E17.5. (B) Six litters with 53 pups from Emb^+/-^ breedings were analyzed at E17.5. The significance of the weight difference between WT and Emb^-/-^ pups were statistically analyzed by Student’s T-test for independent samples. Data are represented as a Spear style box plot. A square shows the mean value. (C) WT and Emb^-/-^ placenta paraffin sections were immunostained with embigin and collagen I antibodies at E11.5 and E17.5. Arrowheads show fetal-derived round nuclei. Scale bars: 100 µm. (D) Representative images of hematoxylin-eosin-stained WT and Emb^-/-^ placentas at E17.5 are shown. The scale bar is 500 µm. Magnifications of the inner edges of the placental labyrinth zones are shown below. The scale bar is 100 µm. (E) Heatmap of differentially expressed genes (log2 of fold change above 0.6 or below -0.6 and Benjamini-Hochberg-corrected p-value < 0.05) in WT (n = 5) and Emb^-/-^ (n = 5) placenta at E17.5 based on RNAseq analysis.

### The maturation of lungs is delayed in embigin deficient embryos

The histological examination of the Emb^-/-^ lungs at E17.5 unveiled the abnormal structure (Fig. 4A): the number and the size of the airways were observed to be significantly smaller (p = 0.00002) when compared to the architecture of the lungs of their WT littermates (Fig. 4B). Further, the relative area of airways of the Emb^-/-^ embryonic lungs at E17.5 was determined to correlate with the size of the Emb^-/-^ embryo strongly (r_s_ = 0.701, p = 0.005, Fig. 4C): the bigger the embryo the more mature were the lungs. However, in Emb^-/-^ embryos, the lung development was defined to be systematically delayed at the canalicular stage at E17.5. This developmental stage is characteristic of the normal mouse lung maturation at E16.5, but it should not be prominent at E17.5.

**Figure 4.**
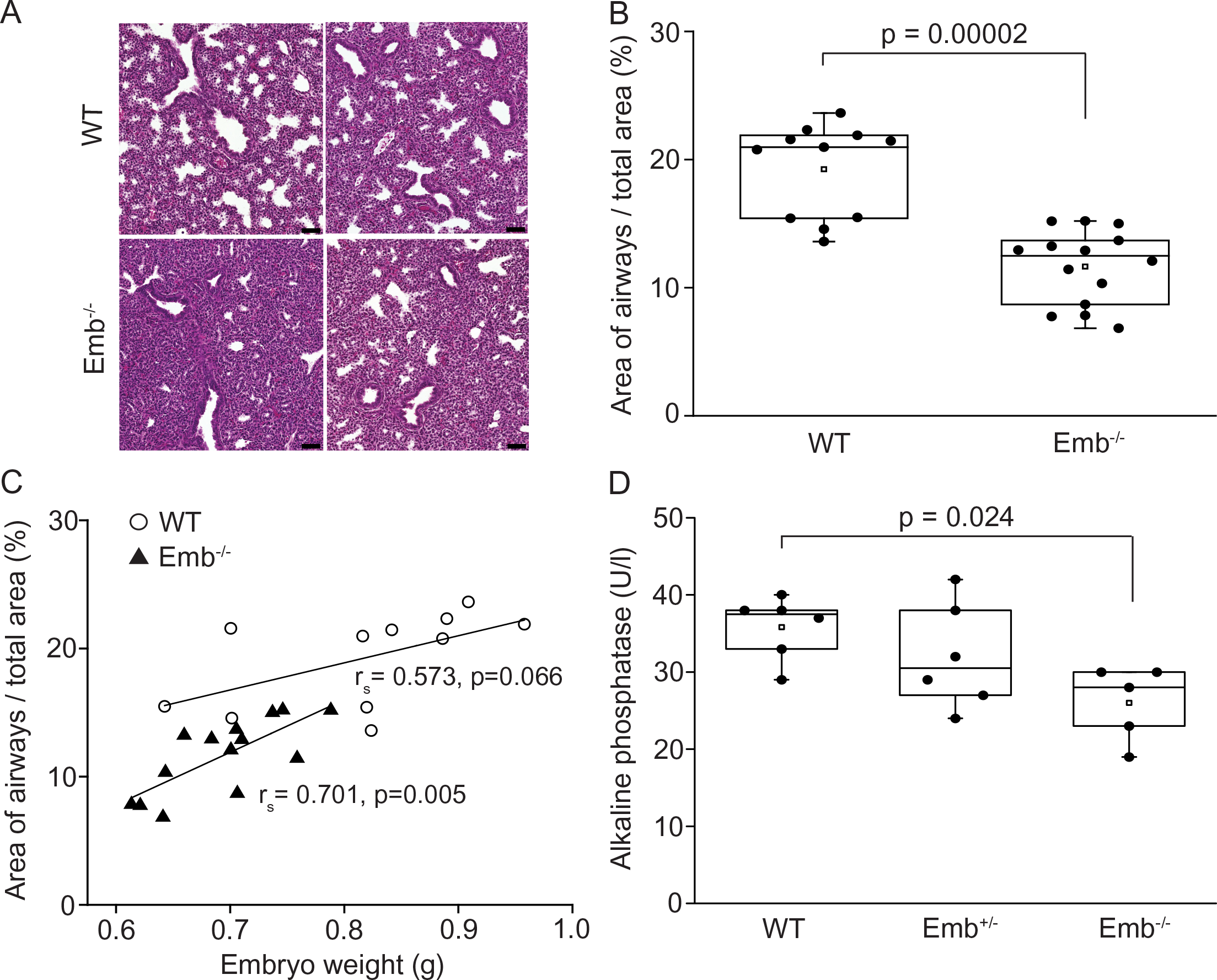
Embigin deficiency delays embryonic lung development. (A) Representative images of hematoxylin-eosin stained lung sections from two WT and two Emb^-/-^ E17.5 littermates are presented. Scale bars: 50 µm. (B) The relative area of airways in the E17.5 lung sections was analyzed by ImageJ/Fiji software (n = 11 for WT and n = 14 for Emb^-/-^). Statistical significance (p = 0.00002) was determined by Mann-Whitney U-test. Data are represented as a Spear style box plot. A square shows the mean value. (C) Correlation between the body weight of the mouse and relative area of airways in the lung sections was studied by calculating Spearman’s rank correlation coefficient (r_s_) and its statistical significance. (D) Alkaline phosphatase activity was determined from the amniotic fluids of WT, Emb^+/-^, and Emb^-/-^ embryos at E17.5 with VetScan Chemistry Analyzer (n = 6 for WT, and n = 5 for Emb^+/-^ and Emb^-/-^). Statistical significance (p = 0.024) was determined by ANOVA followed by Dunnett’s t-test. Data are represented as a Spear style box plot. A square shows the mean value.

Next, the activity of alkaline phosphatase was analyzed in the amniotic fluid at E17.5. Elevated alkaline phosphatase activity at the end of gestation has been shown to indicate increasing fetal lung maturity (Brocklehurst and Wilde, 1980). When the alkaline phosphatase activity of amniotic fluids from WT, Emb^+/-^ and Emb^-/-^ embryos were measured, the average activity was 35.8 U/l for WT, 32 U/l for Emb^+/-^ and 26 U/l for Emb^-/-^ embryos. Thus, the alkaline phosphatase activity in the amniotic fluid of the Emb^-/-^ embryos was significantly lower than the activity detected in WT embryos (p = 0.024). The average alkaline phosphatase activity in the amniotic fluid of Emb^+/-^ embryos neither reached the same level as detected in WT embryos, but the difference was not statistically significant (p = 0.45; Fig. 4D). Not only the activity of the alkaline phosphatase, but also sodium, calcium, and glucose concentrations have been reported to vary in the amniotic fluid during normal pregnancy: their concentrations increase and subsequently decrease as the gestation progresses (Cheung and Brace, 2005). While the alkaline phosphatase activity was significantly lower in the amniotic fluid from Emb^-/-^ embryos at E17.5, the concentrations of sodium, calcium, or glucose did not significantly differ between WT, Emb^+/-^, and Emb^-/-^ embryos (Fig. S5). Only moderate changes observed in the concentrations of these factors indicate that the pregnancies of Emb^-/-^ embryos progress normally. Both the abnormal histological architecture of the fetal lungs and the lower activity of alkaline phosphatase in the amniotic fluids strongly suggest that the maturation of the lungs is delayed in Emb^-/-^ embryos.

To further study the developmental delay, transcriptomes of embryonic lungs at E17.5 were analyzed by RNA sequencing. While the gene expression profiles of the placentas of Emb^-/-^ and WT mice resembled each other (Figs 5A, 3E), in the lungs total 161 genes were differentially expressed at E17.5 between the genotypes (Figs 5A, S4). Particularly genes that participate in cell division were upregulated in the lungs of Emb^-/-^ embryos when compared to WT mice (Fig. 5B), including cell cycle effectors *Ccnf, Cdc6, Cdc45, Cdt1, and Gli1*. Downregulated genes consisted mainly of genes involved in the immune response (Fig. 5B). Additionally, many of these genes are potential transcription factors and growth factors in lung development. For example, *Scgb3a2* is a growth factor in lung promoting both early and late stages of fetal lung development (Kurotani et al., 2008), *Adamts18* is pivotal in airway branching morphogenesis (Lu et al., 2020), and *Hmga2* is required for WNT signaling during lung development (Singh et al., 2014). The data suggest that the lack of embigin causes delays rather than structural defects in lung development. In Emb^-/-^ embryos, the increased expression of cell proliferation-related genes at E17.5 may indicate, that the lungs execute an earlier stage of differentiation compared to WTs. The crucial function of embigin especially during the earlier stages of development is supported by the wave-like changes in its expression profile: the protein is prominent in the early days of development (Fig. 1A), only weakly expressed in embryonic lungs at E13.5 (Fig. S1A) and below detection level in lungs at E17.5. At mRNA level, embigin is clearly present in lungs at E17.5 (Fig. S4), and embigin protein expression rises again after birth (Fig. 1C). Furthermore, the lifespan or the histological architecture of lungs are not affected in Emb^-/-^ mice that survive to adulthood (Fig. S3). Thus, the results indicate that embigin deficiency directly affects lung development, which explains well the detected high perinatal mortality of Emb^-/-^ mice.

**Figure 5.**
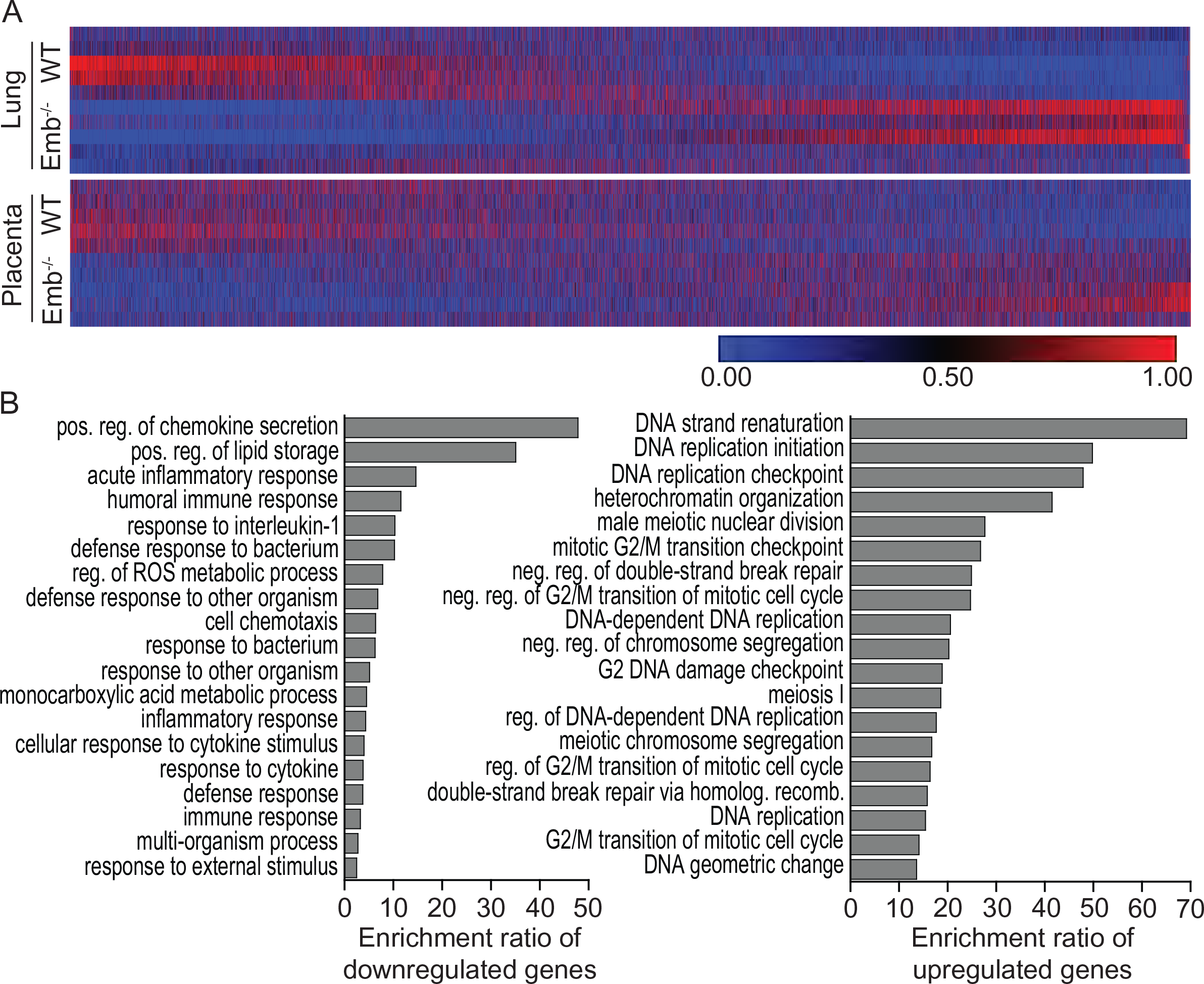
Embigin depletion leads to compromised lung function. (A) Heatmap of all expressed genes in WT (n = 5) and Emb^-/-^ (n = 5) lung and placenta at E17.5 based on RNAseq analysis. The columns (genes) have been sorted from smallest to largest fold change value of Emb^-/-^ vs. WT comparison and visualized by Morpheus. (B) Differentially expressed genes in WT (n = 5) vs. Emb^-/-^ embryonic lungs (n = 5) at E17.5 were analyzed from RNA sequencing data by WebGestalt. Top19 of the most enriched biological processes of those with FDR < 0.05 and ≥ 3 overlapping genes are shown.

In conclusion, our study provides the first characterization of embigin knockout mice. We demonstrate that embigin is normally expressed in the adult mouse lung, kidney, skin, epididymis, and testis. Furthermore, the study emphasizes the role of embigin during mouse embryonic development: embigin deficiency leads not only to growth retardation of the Emb^-/-^ embryo but also to high, 72%, mainly neonatal mortality. Though embigin is expressed in the placenta, no such signs of placental dysfunction were detected that could explain the delay in embryonic growth of the Emb^-/-^ mice. Instead, the data suggest that the increased lethality of Emb^-/-^ mice was primarily due to developmental delays, rather than structural defects, in embryonic lungs.

## Discussion

Embigin has remained a less studied member of the basigin subgroup in the immunoglobulin superfamily. Here, we describe embigin expression and its role in mouse development using embigin deficient mice. Our data reveal that embigin is a widely expressed protein during entire mouse embryonic development. In adult mice, embigin expression is restricted mainly to epithelial cells lining tubular structures in the lung, kidney, and epididymis, in addition to skin and testis. Given that 72% of Emb^-/-^ offsprings are lost latest in the neonatal period and the Emb^-/-^ deficient fetuses are typically smaller with a delay in lung maturation, embigin can be considered as a pivotal protein for mouse development.

Our findings indicate that embigin is an abundantly expressed protein in the developing mouse embryo during the first half of gestation. This observation is in agreement with the previous data gained from the analyses performed at mRNA level: embigin mRNA has been described to be moderately expressed at mouse embryonic days E5 - E6 and strongly present at E7 - E9. However, the embigin expression was reported to disappear after E9, and only a weak expression of embigin mRNA was observed in adult animals (Fan et al., 1998;Huang et al., 1990). In line with the mRNA studies, we observed a decrease in the embigin protein level after E10.5, yet we could detect embigin protein throughout the embryonic period. In adult mice, we observed the embigin expression to be restricted to specific organs, i.e., the epithelial cells of the lung and kidney, sebaceous glands in the skin, as well as in epididymis and testis. Tabula Muris, a compendium of single-cell transcriptome data from the 3-month-old mice, supports our findings showing that embigin is present, for instance, in the epithelial cells of lungs and in the epithelial cells of collecting duct in the kidney (The Tabula Muris Consortium, 2018). In addition to the restricted expression pattern of the embigin in adult mice, we observed it to be differentially glycosylated in each tissue. As in the case of basigin, this might propose the tissue-specific function of the protein (Bai et al., 2014;Tang et al., 2004).

Embigin deficiency compromised the viability of Emb^-/-^ offspring. Despite we observed the high embigin expression during the first embryonic days, embigin deficiency did not increase the mortality of Emb^-/-^ mice on this particular phase of gestation. Instead, the fetal resorption among Emb^-/-^ embryos started to increase slightly after embryonic day E8.5. However, the highest occurrence of loss of Emb^-/-^ offspring was in the neonatal period. When compared to the phenotype of basigin null animals (Bsg^-/-^), the presence of embigin seems to be less critical than basigin during embryonic development: the majority, about 70%, of Bsg^-/-^ embryos die during the early stages of embryonic development. Consequently, basigin has been suggested to be involved in intercellular recognition during implantation and/or early post-implantation stages (Igakura et al., 1998). In addition to the higher survival rate during the embryonic period, Emb^-/-^ mice also survived better (28%) than Bsg^-/-^ mice (14%) after birth. Furthermore, while half of the survived Bsg^-/-^ mice have been observed to die in interstitial pneumonia during the first month after birth (Igakura et al., 1998), the Emb^-/-^ mice, that successfully passed the early neonatal period, seemed to have an unaffected lifespan. For an unknown reason, the survival of Emb^-/-^ females was slightly better than males. We also show that embigin null animals were able to produce offspring, while Bsg^-/-^ males have been reported to be sterile, and also Bsg^-/-^ females have been detected with fertility problems (Igakura et al., 1998;Kuno et al., 1998). The observed phenotype of Emb^-/-^ mice is also very different when compared to neuroplastin deficient animals. For example, Np65 null mice have been reported to show deviant behavior in cognitive tests (Amuti et al., 2016), and they suffer from the loss of hearing (Carrott et al., 2016). Thus, it can be summarized that all three members of the basigin family have separate functions.

The mortality among embigin null embryos started to rise after day E8.5. During the pregnancy, the Emb^-/-^ embryos were also significantly smaller than their WT littermates. Since the placenta is pivotal for normal fetal growth and development, the dysfunction of the placenta might explain the delay in embryonic growth and development. Indeed, we detected fetal-derived embigin to be an abundant protein in the labyrinthine layer of the placenta, and its expression seemed to increase toward the later stages of gestation. However, only minor changes were observed when the gene expression patterns of the placentas of Emb^-/-^ and WT mice were compared,. Neither the placental architecture was affected in Emb^-/-^ embryos. The data suggest that placental dysfunction may not be the primary cause for the growth delay of Emb^-/-^ embryos during the second half of gestation.

Because we did observe neither typical dysmorphological features in the placenta of Emb^-/-^ fetuses nor correlation with typical fetal organ defects caused by the impaired placenta (Perez-Garcia et al., 2018), the primary reason for the loss of Emb^-/-^ offspring might reside in the embryo itself. We did not detect any apparent defects in other major tissues of Emb^-/-^ embryos, however, the significantly delayed morphogenesis in the fetal lungs was characterized. Histologically, lung development and maturation has been divided into four stages: pseudoglandular, canalicular, terminal saccular, and alveolar (Warburton et al., 2010). The Emb^-/-^ lung development at E17.5 was observed to be delayed at the canalicular stage which is characteristic of the normal mouse lung maturation at E16.5. As expected, the WT lung showed typical lung morphology for the terminal saccular stage at E17.5. Since the elevated level of alkaline phosphatase activity in the amniotic fluid at the end of gestation has been shown to correlate with fetal lung maturity (Brocklehurst and Wilde, 1980), the low alkaline phosphatase activity detected in the amniotic fluid of Emb^-/-^ further confirmed the delay in the lung development of the Emb^-/-^ embryos. Maturity-related increase in alkaline phosphatase activity has also been reported in the epithelial cells that line the airway cavities in the embryonic murine lungs (Sasaki and Kahn, 2014). Based on the function of alkaline phosphatase, glucocorticoids, which increase the activity of alkaline phosphatase in some conditions, have been used for decades to induce the maturation of the preterm fetal lungs (Green et al., 1990;Grier and Halliday, 2004). On this basis, the lungs of the Emb^-/-^ mice were suggested to be underdeveloped at the stage of birth, which explains the remarkably increased neonatal mortality of the affected animals.

Lung development is controlled by various transcription factors and growth factors, and some of the upregulated or downregulated genes we observed in E17.5 Emb^-/-^ mice can be directly linked to lung development. We also discovered that several genes involved in cell proliferation were upregulated in Emb^-/-^ mice implicating the earlier stage of lung development compared to WT mice. During the canalicular stage, a massive increase in the cell mass occurs during the formation of the most distal airways (Warburton et al., 2010) airways. Overall, we suggest that embigin causes primarily a developmental delay in lung maturation rather than structural defects. Noteworthy, at the same developmental stage, E9, where we observed the first losses of Emb^-/-^ embryos, the organogenesis of lungs begins (Warburton et al., 2010). Embigin expression was detected to be highest at the early developmental days, gradually disappearing from the embryonic lungs only to be increased again after birth. Given that embigin deficient mice that survive do not display changes in their lifespan or in the histological lung architecture, the critical function of embigin can be placed on the early days of development.

In summary, our results indicate that embigin is a critical protein for the proper morphogenesis of the mouse lungs during development. Delayed maturation of embryonic lungs explains why the majority of Emb^-/-^ mice are lost during the neonatal period. However, given the abundant expression of embigin and the nature of other basigin family members as multifunctional proteins, it is possible that Emb^-/-^ mice have several defects that simultaneously contribute to the Emb^-/-^ knockout phenotype. To conclude, our results indicate that abundantly expressed embigin is a vital protein for overall embryonic development and for lung maturation that explains the high mortality of Emb^-/-^ embryos.

## Materials and methods

### REAGENTS

#### Cell lines

G4 embryonic stem cells derived from 129S6/SvEvTac x C57BL/6NCrl mice (Mutant Mouse Resource & Research Center (MMRRC)) were cultured on neomycin-resistant primary embryonic fibroblast (Neo-resistant MEF feeder cells, Applied StemCell) feeder layer in KnockOut DMEM medium (Gibco, Thermo Fisher Scientific) supplemented with 10% ES screened fetal bovine serum, heat-inactivated (Cytiva). The cell lines were cultured at 37°C in a humidified atmosphere with 5% CO_2_.

#### Antibodies

Following antibodies were used in our studies: Embigin Monoclonal Antibody, clone G7.43.1, 14-5839-81, Lot#4343173, eBioscience, Thermo Fisher Scientific (Western blotting 1:1000, whole-mount and immunofluorescence 1:200); Monoclonal Anti-β-Tubulin I antibody produced in mouse, clone SAP.4G, T7816, Lot#068M4850V, Sigma-Aldrich (Western blotting 1:20 000); Anti-α smooth muscle Actin (α-SMA) antibody [1A4] (Alexa Fluor 488), ab184675, Lot#GR316286-7, Abcam (whole-mount: 1:250); Collagen I Antibody, NB600-408, Lot#41476, Novus Biologicals (immunofluorescence 1:300); IRDye secondary antibodies, LI-COR Biosciences (Western blotting 1:15 000); and Alexa Fluor secondary antibodies, Thermo Fisher Scientific (whole-mount and immunofluorescence 1:400).

### ANIMAL MODELS

C57BL/N6 mice (*Mus musculus*, Charles River Laboratories, Willmington, MA) and the generated embigin knockout mice (collaboration with Turku Center for Disease Modeling) were maintained in Central Animal Laboratory at the University of Turku, Finland. All animal experiments were formally reviewed and approved by the Ethical Committee for Animal Experimentation in Finland, complying with international guidelines on the care and use of laboratory animals. The mouse embryos were examined between embryonic days E8.5-E17.5 and pups at postnatal days P0-P3. Both male and female adult mice were studied at the age of 2, 4, or 6 months.

### METHOD DETAILS

#### RNA sequencing

Five E17.5 WT and Emb^-/-^ placentas and lungs were dissected, and the RNA was isolated as described. The libraries were prepared from 300 ng of RNA from each sample using TruSeq Stranded mRNA HT Kit and TruSeq Stranded mRNA Sample Preparation protocol 15031047 (Illumina). Sequencing was performed with NovaSeq 6000 SP Sequencing System (Illumina) using paired-end sequencing chemistry and 2 x 50 bp read length. The reads obtained from the instrument were base called using bcl2fastq2 conversion software. Raw data were obtained as fastq-files, which were uploaded to Chipster (Kallio et al., 2011). The reads were aligned against the reference genome (Mus musculus GRCm38.95, available in Chipster) using STAR, version 2.7.3 (Dobin et al., 2013). The reads associated with each gene were counted using the HTSeq package, version 0.12.4 (Anders et al., 2015).

The edgeR R/Bioconductor package (Robinson et al., 2010) was used to normalize gene-wise read counts by TMM normalization method and to perform statistical tests between groups. The results were filtered to have a minimum of 50 reads per gene in at least one sample. The gene was determined as differentially expressed if the following conditions were met: log2 of fold change value was above 0.6 or below -0.6 and Benjamini-Hochberg-corrected p-value less than 0.05. WebGestalt, http://www.webgestalt.org/ (Liao, Y. et al., 2019), was used to perform over-representation analysis of differentially expressed genes against biological process gene ontology. Morpheus (https://software.broadinstitute.org/morpheus) was used to generate heatmaps of the differentially expressed genes. For generating heatmaps by Morpheus (https://software.broadinstitute.org/morpheus), the raw counts were first transformed by using deseq-transform function of DEseq2 package (Love et al., 2014).

#### Generation of embigin deficient (Emb^-/-^) mice

Emb^-/-^ mice were generated in collaboration with Turku Center for Disease Modeling. First, a targeting vector for Emb gene, HTGR06008_A_1_E08 from The European Conditional Mouse Mutagenesis Program, was linearized with AsiSI restriction enzyme (R0630S, NEB). Construct was then transfected by electroporation into G4 embryonic stem cells derived from 129S6/SvEvTac x C57BL/6NCrl mice and cultured on neomycin-resistant primary embryonic fibroblast feeder layer for 7-9 days. To ensure the occurrence of the correct homologous recombination, positive ES cell clones were screened by PCR and sequencing. ES cells were injected into C57BL/N6 mouse blastocysts to generate chimeric mice. Germline transmission was achieved by cross-breeding male chimeras with C57BL/N6 females.

#### Timed matings and genotype determination

In timed matings, the day of vaginal plug appearance was considered as embryonic day 0.5 (E0.5). To analyze the survival of the Emb^-/-^ embryos, the genotypes from 203 pups from 25 litters and 25 Emb^+/-^ breedings were determined between embryonic days E8.5-E17.5. Furthermore, the genotypes of 100 pups from 17 litters from six different Emb^+/-^ breedings were analyzed between postnatal days P0-P3; and the genotypes from 284 pups from 40 different litters from Emb^+/-^ breedings were analyzed at P14-P21. Genomic DNA was extracted with gDNA Nuclespin tissue kit (Macherey-Nagel) and the genotypes of the mice were determined from genomic DNA by using a PCR primer pair 1 (5’-TAAGTCTCTTGTTTGCTGTG-3’; 5’-CACAACGGGTTCTTCTGTTAGTCC-3’) to detect embigin knockout allele and a PCR primer pair 2 (5’-ACCCTTAAGTGCATGAACAAAA-3’; 5’-GGGTTCCTTGGCATTGTTACTAA-3’) to detect embigin WT allele. DreamTaq polymerase (Thermo Fisher Scientific) was used according to the manufacturer’s instructions using following reaction settings: 95 °C, 2 min; 35 x [95 °C, 30 s; 50 °C, 30 s; 72 °C, 1 min]; 72 °C, 10 min.

#### Emb^-/-^ mice fertility

The fertility of Emb^-/-^ mice was studied with ten Emb^-/-^ breedings. The crossings were followed until the pups were genotyped at the age of P14-P21. Viable pups and average litter size were examined.

#### Size of embryos at E17.5

The body weights of the embryos from Emb^+/-^ breedings were weighted at E17.5. Six litters with 53 pups were analyzed in total.

#### Histological staining

Male and female WT and Emb^-/-^ were examined at the age of 2, 4, and 6 months. Three to four mice were included in each independent study group. The mice and the specific organs, heart, lung, liver, spleen, kidney, epididymis, testis, and ovary, were weighted, and in addition to the samples of skin, small intestine and adrenal glands were prepared for histological analysis. Lungs were additionally analyzed from 11 WT and 14 Emb^-/-^ mice from six E17.5 Emb^+/-^ breedings. Placentas were analyzed at E17.5. Formalin-fixed samples were fixed in paraffin and 4 μm sections were cut using an RM2255 microtome (Leica) and immobilized to adhesion slides (SuperFrost Plus, Thermo Fisher Scientific) overnight at 37 °C. The sections were deparaffinized, rehydrated, and stained with conventional hematoxylin and eosin (HE), imaged with Pannoramic 250 Flash III slide scanner (3D Histech), and analyzed with CaseViewer program (3D Histech). In the case of HE-stained histological lung section from E17.5 embryos, three images per organ section at 20x magnification were selected with the CaseViewer program. The relative area of airways in the lung section images was analyzed with ImageJ/Fiji (Schindelin et al., 2012).

#### Western blotting

The expression of embigin in the lung, kidney, skin, heart, liver, spleen, small intestine, adrenal gland, epididymis, testis, and ovary, of WT mice and the kidneys of Emb^-/-^ mice at the age of four months was analyzed with Western blotting. Protein samples were extracted from the organs with the NucleoSpin RNA/Protein kit (Macherey-Nagel). Macherey-Nagel Bead Tubes Type F was used in tissue homogenization. Protein concentrations were measured with Pierce 660nm Protein Assay (Thermo Scientific), and 10 μg of protein were loaded on 4–20% FastGene SDS-PAGE gradient gels (Nippon Genetics). Embigin and β-tubulin (diluted in 5% milk and 0.1% Tween-20 in TBS) were stained in the membrane for 2 hours at RT. IRDye secondary antibodies and Odyssey CLx imager (LI-COR Biosciences) were used for signal detection.

#### Whole-mount fluorescent immunohistochemistry

The embryonic embigin expression was analyzed using a whole-mount immunostaining technique as described previously (Yokomizo et al., 2012). However, PBS-MT solution was replaced with PBS-BSA-T (1% (w/v) bovine serum albumin (BSA) and 0.4% (v/v) Triton X-100 in PBS). 1% (v/v) normal mouse serum (10410, Invitrogen) and 0.5% (v/v) fetal calf serum solution (PromoCell) in PBS-BSA-T was used as blocking solution. WT embryos at E8.5, E9.5, and E10.5 and Emb^-/-^ embryos at E9.5 were stained with embigin and α-SMA antibodies. In the negative control for embigin, a secondary antibody only was applied. α-SMA was used as a positive control. The embryos were imaged with LSM 880 confocal microscope (Zeiss) using Plan-Apochromat 20x/0.8 M27 objective for E8.5 embryos and Plan-Apochromat 10x/0.3 M27 for E9.5 and E10.5 embryos. Image stacking, background subtractions, linear brightness, and contrast adjustments were performed with Zeiss ZEN blue software and Imaris (Bitplane).

#### Immunofluorescence

Embigin expression was analyzed in the paraffin sections of the whole embryo at E13.5 in addition to the paraffin sections of lung and kidney at E17.5 and postnatal day 3 (P3). Placental paraffin sections were examined at E11.5 and E17.5. The expression was further examined in the paraffin sections of the lung, kidney, skin, heart, liver, spleen, small intestine, adrenal gland, epididymis, testis, and ovary from three four-month-old male and female. 4 μm sections were cut and immobilized to adhesion slides as mentioned earlier. The sections were deparaffinized and rehydrated. The antigen retrieval was achieved with 3 min proteinase K treatment (S3020, Agilent), and the sections were washed in PBS. The samples were blocked with 1% (v/v) BSA in PBS for 1 h at RT, and stained with antibodies against embigin and collagen I in blocking buffer o/n at 4 °C. The samples were washed with PBS and incubated with Alexa Fluor secondary antibodies in blocking buffer for 1 h at RT. The sections were washed with PBS and nuclei were labeled with Hoechst 33342 (1:5000 in PBS, Thermo Fisher Scientific) for 10 min at RT. The sections were rinsed in PBS and finally in dH2O and mounted in Mowiol (Calbiochem) containing 25 mg/ml DABCO anti-fading reagent (Sigma). The samples were imaged with LSM 880 confocal microscope (Zeiss) using a Plan-Apochromat 20x/0.8 M27 objective. Image stacking, background subtractions, linear brightness, and contrast adjustments were performed with ImageJ/Fiji software.

#### Alkaline phosphatase in amniotic fluid

Amniotic fluids were collected from six WT, five Emb^-/-^, and five Emb^+/-^ embryos at E17.5 and analyzed with VetScan Comprehensive Diagnostic Profile reagent rotor (Abaxis) used with the VetScan VS2 Chemistry Analyzer (Abaxis). Alkaline phosphatase activity (U/l) and the molar concentrations (mmol/l) of sodium, calcium, and glucose were determined.

### STATISTICAL ANALYSIS

IBM SPSS Statistics software (version 25, IBM) was used for all statistical analyses. Correlation between the frequency of each genotype and embryonic stage, as well as the correlation between the body weight of the mouse and relative area of airways in the lung sections, were analyzed with the Spearman’s rank correlation test. Statistical significance of the differences of the relative area of the airways between WT and Emb^-/-^ mice was determined with Mann-Whitney U-test. The significance of the weight difference between WT and Emb^-/-^ pups was analyzed statistically by Student’s T-test for independent samples. When amniotic fluids were analyzed, the normality of the data was checked with the Shapiro-Wilk test and Levene’s test was used to determine the equal variances. Normally distributed alkaline phosphatase data were analyzed using ANOVA (p = 0.039) followed by Dunnett’s two-sided t-test. Normally distributed Ca^2+^ was analyzed with ANOVA (p = 0.219) only. A nonparametric alternative for ANOVA, Kruskal-Wallis H-test with exact p-value, was used for glucose and sodium data. A p-value of less than 0.05 was considered statistically significant. Statistical details of the experiments can be found in the figures and in the figure legends.

## Acknowledgements

The authors would like to thank the personnel of Turku Center for Disease Modeling for assistance. We are grateful for the personnel in the Central Animal Laboratory of the University of Turku. Maria Tuominen is acknowledged for excellent technical assistance.

## Competing interests

No competing interests declared.

## Funding

This study has been financially supported by grants from the Academy of Finland [259769 to JH], Sigrid Juséliuksen Foundation [to JH], Cancer Society of Finland [to JH], and The Finnish Foundation for Cardiovascular Research [to JH].

## Data availability

RNA-seq data have been deposited in the ArrayExpress database at EMBL-EBI (www.ebi.ac.uk/arrayexpress) under accession number E-MTAB-10641 (Username: Reviewer_E-MTAB-10641, Password: bdvimnnp).

## SUPPLEMENTAL DATA

**Figure S1.**
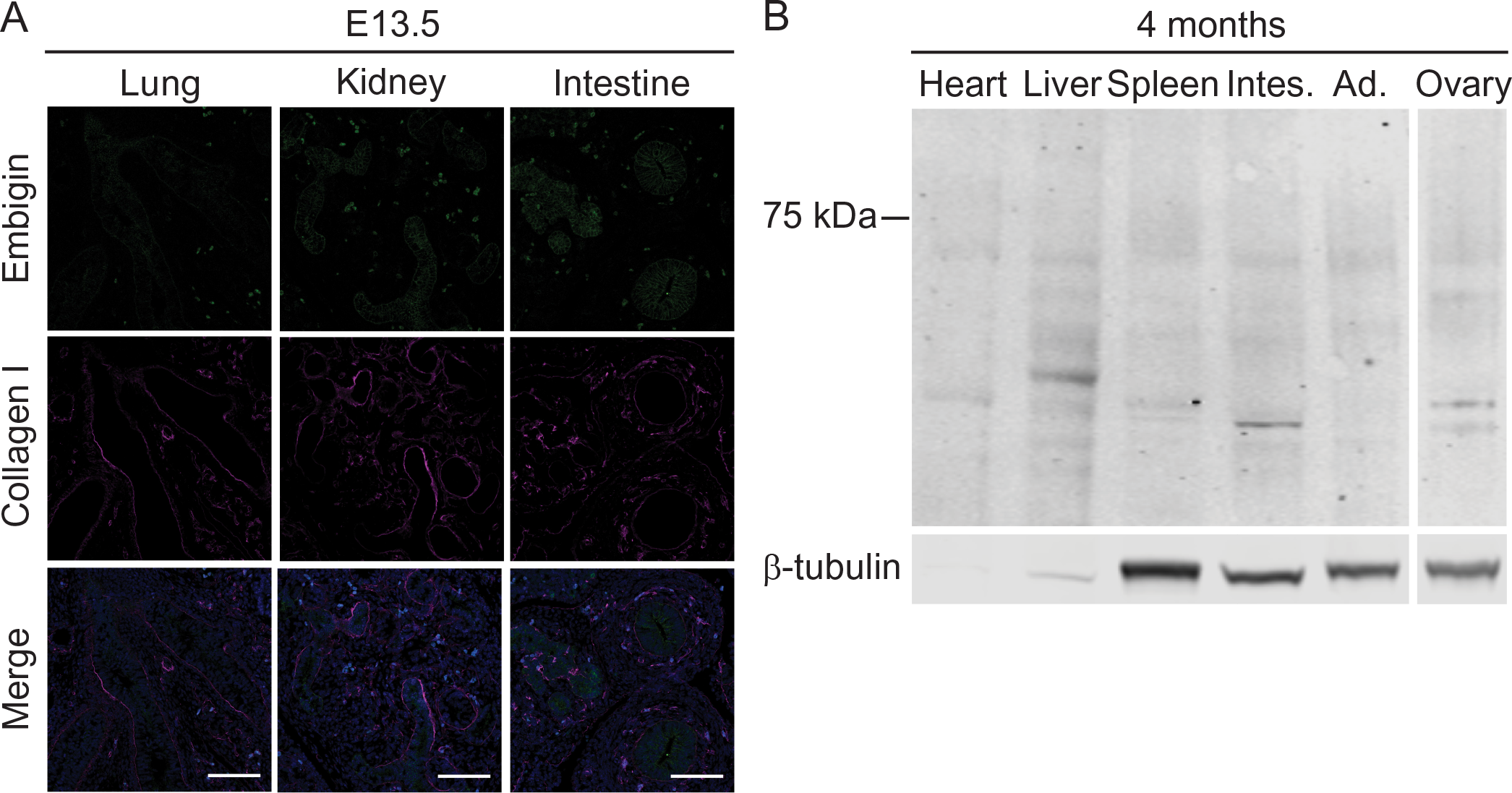
Embigin protein has a tissue-specific expression pattern. (A) The paraffin sections of embryos at E13.5 were immunostained with embigin and collagen I antibodies. Embigin expression in the lung, kidney and intestine is shown. Scale bars: 100 µm. (B) Embigin expression in protein samples extracted from four-month-old WT mouse heart, liver, spleen, small intestine (Intes.), adrenal gland (Ad.), and ovary tissues were analyzed by Western blotting. β-tubulin was used as a positive control. The blot image was cut to show only embigin negative organs.

**Figure S2.**
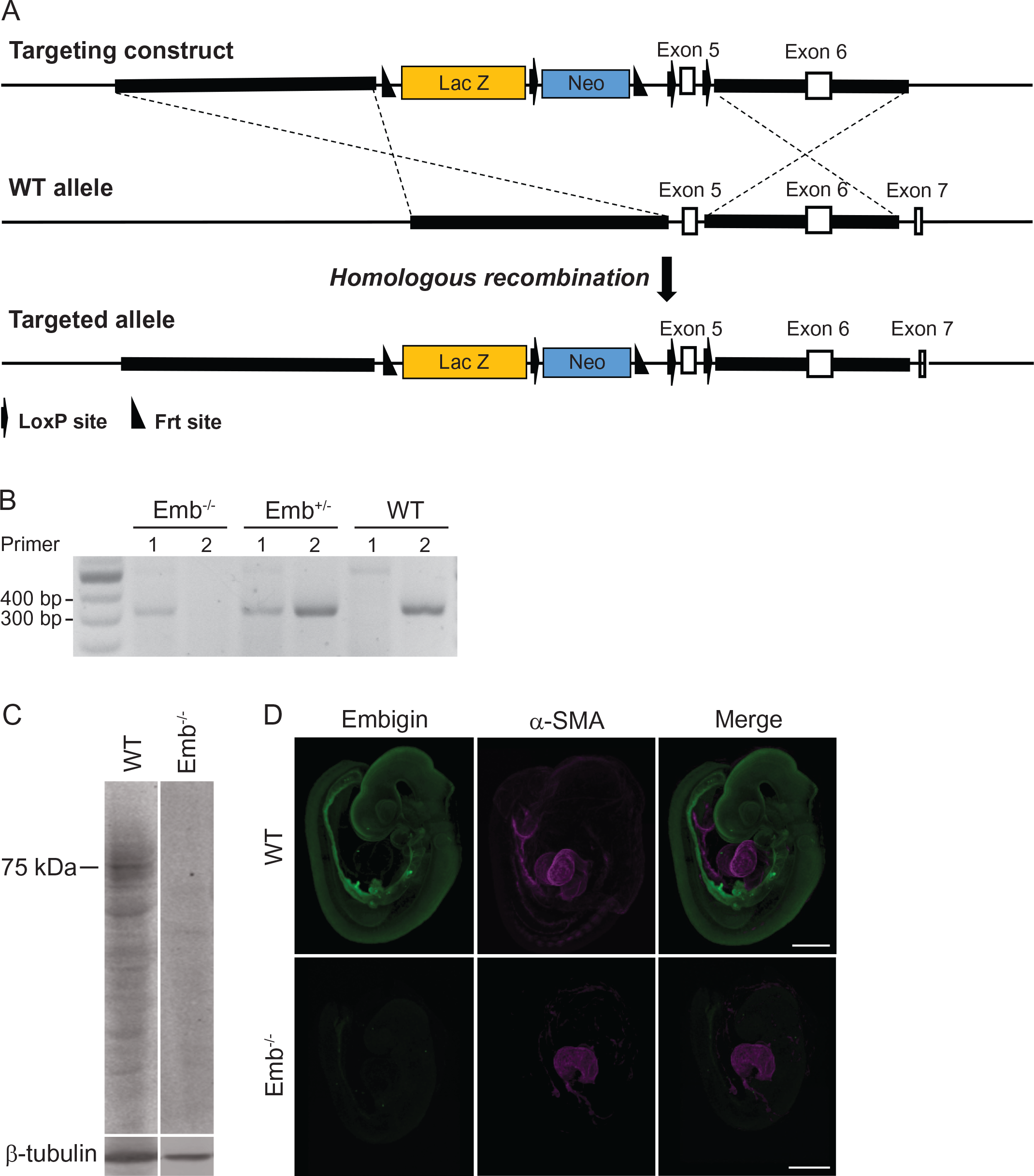
Generation of embigin knockout mice. (A) The knockout-first strategy was used to create knockout alleles. In the schematic presentation, the structure of the targeting construct is presented at the top, the wild type embigin allele with the coding exons 5, 6, and 7 in the middle and the homologously mutated allele below. Homologous sequences in the targeting construct are presented with a bold line; arrows represent loxP sites and triangles Frt sites. (B) The genotypes of the mice were determined from genomic DNA by using PCR primer pair 1 to detect embigin knockout allele (354 bp) and PCR primer pair 2 to detect embigin WT allele (349 bp). (C) The expression of embigin in the kidney tissue of WT and Emb^-/-^ mice at the age of four months was analyzed by Western blotting. β-tubulin was used as a control. (D) The expression of embigin in the embryo was analyzed using the whole-mount immunostaining technique. WT and Emb^-/-^ embryos at stage E9.5 were stained with embigin and α-SMA antibodies. α-SMA was used as a positive control. Scale bars: 500 µm.

**Figure S3.**
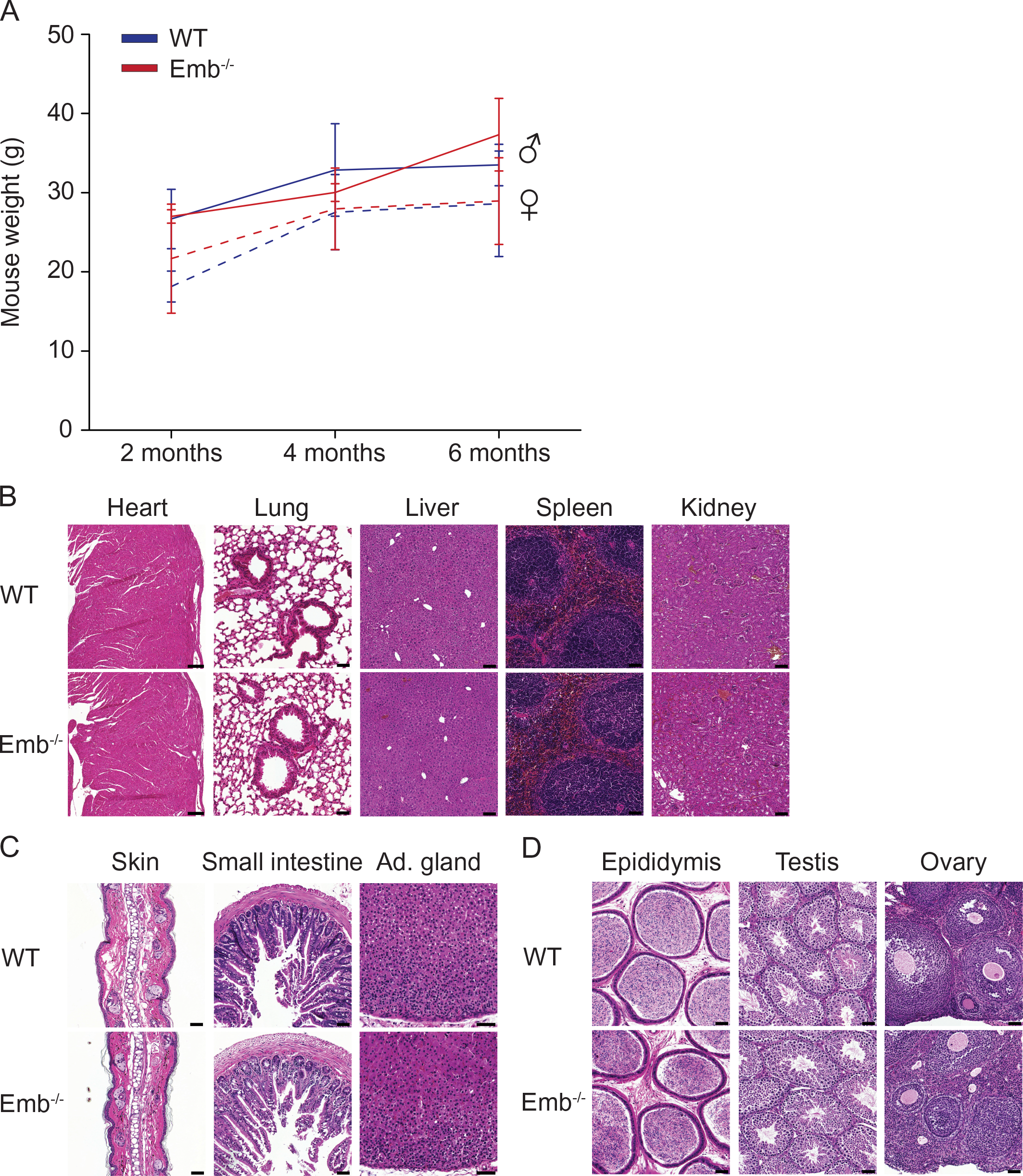
Emb^-/-^ mice that survive into adulthood do not differ from WT mice. (A-D) Male and female WT and Emb^-/-^ were examined at the age of 2, 4, or 6 months. Three mice were included in each independent study group. At each time point, the body weights of the mice were measured (A) and specific organs, heart, lung, liver, spleen, kidney, skin, small intestine, adrenal gland (Ad. gland), epididymis, testis, and ovary, were collected for histological analysis. Representative images of hematoxylin-eosin stained sections from male organs and female ovary at four months of age are shown (B-D). Scale bars: 200 µm (heart), 100 µm (liver and kidney), and 50 µm (other organs).

**Figure S4.**
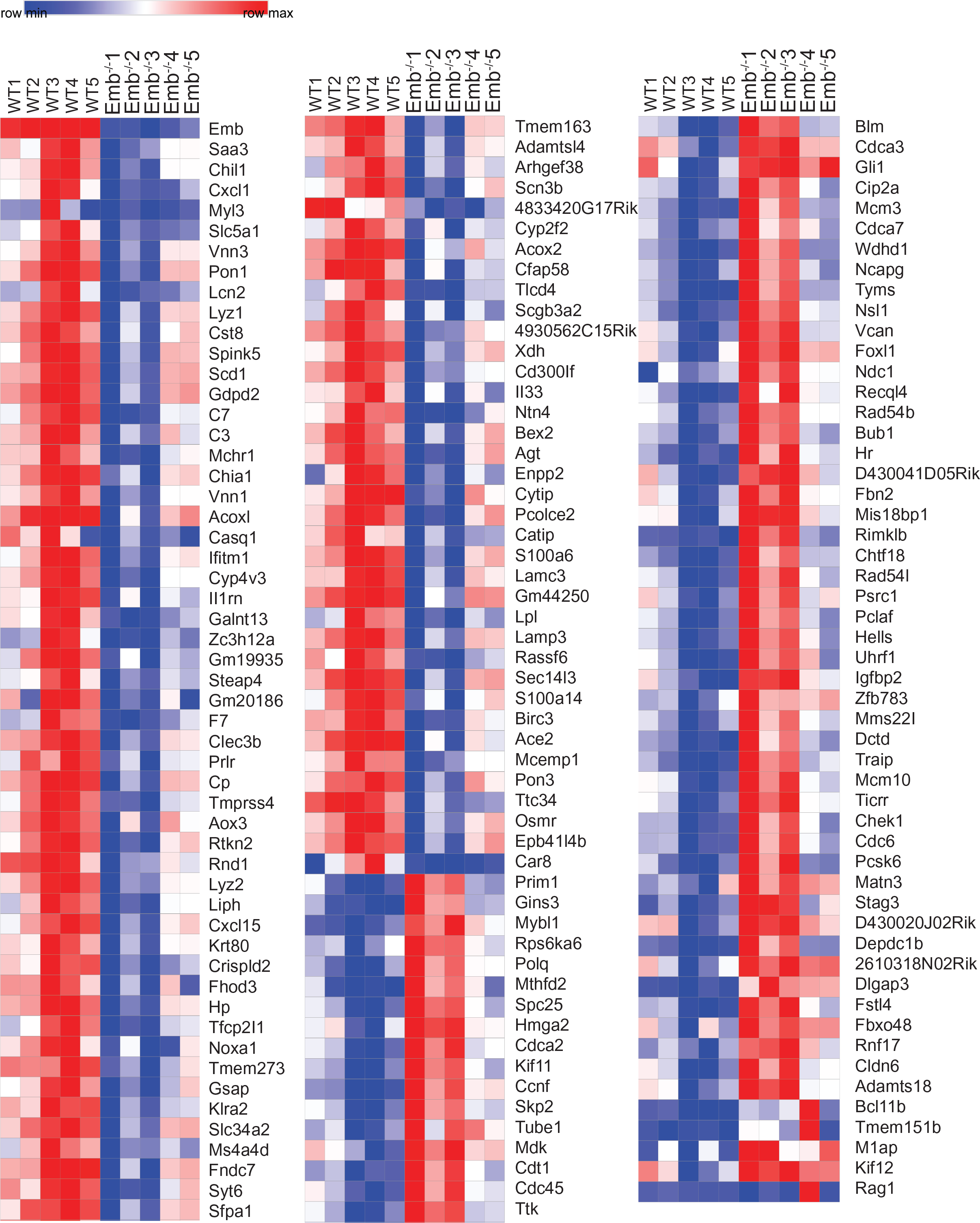
Several genes are differentially regulated in Emb^-/-^ lungs at E17.5. Heatmap of differentially expressed genes (log2 of fold change above 0.6 or below -0.6 and Benjamini-Hochberg-corrected p-value < 0.05) in WT (n= 5) and Emb^-/-^ (n= 5) lungs at E17.5 based on RNAseq analysis.

**Figure S5.**
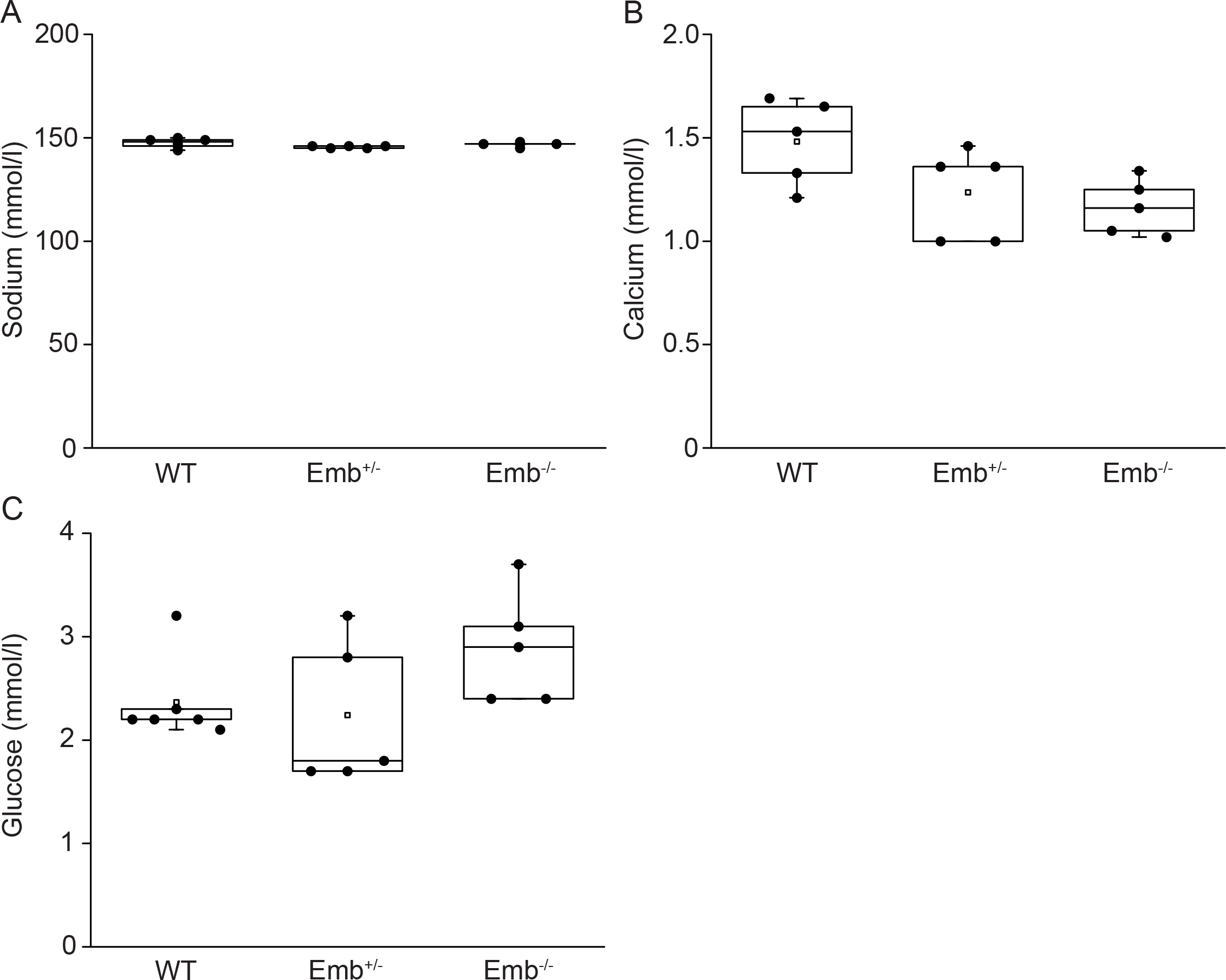
Sodium, calcium, and glucose levels do not vary in amniotic fluids at E17.5. Molar concentrations (mmol/l) of sodium (A), calcium (B), and glucose (C) were determined from WT, Emb^+/-^, and Emb^-/-^ embryo amniotic fluids at E17.5 with VetScan Chemistry Analyzer (n = 6 for WT, and n = 5 for Emb^+/-^ and Emb^-/-^ mice). There were no statistically significant differences between group means determined by one-way ANOVA for calcium or by Kruskal-Wallis H-test for sodium and glucose. Data are represented as Spear style box plots. A square shows the mean value.

**Table S1.** ̶ Emb^-/-^ mice organ weights do not differ from WT mice. Male and female WT and Emb^-/-^ heart, lung, liver, spleen, kidneys, epididymides, and testes were weighed at the age of 2, 4, or 6 months. Three to four mice were included in each independent study group. Relative organ weights were calculated by organ weight/body weight; n = number of organs analyzed.

| Organ | Male 2 mo |  | Male 4 mo |  | Male 6 mo |  |
| --- | --- | --- | --- | --- | --- | --- |
|  | WT<br>(n = 3) | Emb<sup>-/-</sup><br>(n = 3) | WT<br>(n = 3) | Emb<sup>-/-</sup><br>(n = 3) | WT<br>(n = 3) | Emb<sup>-/-</sup><br>(n = 4) |
| Heart | 0.64±0.04 | 0.78±0.10 | 0.67±0.12 | 0.75±0.06 | 0.58±0.07 | 0.71±0.11 |
| Lung | 0.66±0.02 | 0.90±0.36 | 0.55±0.13 | 0.63±0.07 | 0.50±0.03 | 0.66±0.19 |
| Liver | 5.43±0.40 | 5.47±0.90 | 4.63±0.70 | 5.07±1.03 | 5.03±0.69 | 5.59±0.81 |
| Spleen | 0.31±0.06 | 0.44±0.11 | 0.32±0.04 | 0.49±0.21 | 0.22±0.05 | 0.39±0.15 |
| Kidney | 0.82±0.05 | 0.86±0.06 | 0.76±0.09 | 0.96±0.11 | 0.74±0.13 | 0.81±0.21 |
| Epididymis | 0.13±0.04 | 0.12±0.04 | 0.09±0.01 | 0.10±0.02 | 0.10±0.01 | 0.11±0.02 |
| Testis | 0.26±0.07 | 0.26±0.02 | 0.21±0.03 | 0.29±0.02 | 0.21±0.04 | 0.28±0.02 |

| Organ | Female 2 mo |  | Female 4 mo |  | Female 6 mo |  |
| --- | --- | --- | --- | --- | --- | --- |
|  | WT<br>(n = 3) | Emb<sup>-/-</sup><br>(n = 3) | WT<br>(n = 3) | Emb<sup>-/-</sup><br>(n = 3) | WT<br>(n = 3) | Emb<sup>-/-</sup><br>(n = 3) |
| Heart | 0.61±0.11 | 0.60±0.05 | 0.47±0.07 | 0.56±0.13 | 0.54±0.08 | 0.73±0.19 |
| Lung | 0.87±0.09 | 0.96±0.35 | 0.61±0.11 | 0.61±0.21 | 0.55±0.08 | 0.62±0.15 |
| Liver | 5.69±1.36 | 6.05±0.48 | 5.19±0.28 | 4.75±1.07 | 4.73±1.32 | 5.25±0.52 |
| Spleen | 0.45±0.08 | 0.43±0.08 | 0.38±0.10 | 0.49±0.17 | 0.45±0.10 | 0.36±0.05 |
| Kidney | 0.70±0.04 | 0.69±0.09 | 0.55±0.09 | 0.62±0.11 | 0.59±0.11 | 0.66±0.20 |
| Ovary | 0.022±0.007 | 0.026±0.009 | 0.021±0.006 | 0.027±0.009 | 0.015±0.002 | 0.018±0.004 |

**Relative organ weight = organ weight / body weight \* 100 %**

## Notes

### Competing Interest Statement

The authors have declared no competing interest.

